# Disruption of GABA-Regulated Network Resilience in Human Cerebral Organoids Leads to Fragmented Small-World States and Reduced Connectivity

**DOI:** 10.1101/2025.06.11.659143

**Authors:** Sriram Ayalavarapu, Natalie Smith, Zane R. Lybrand

## Abstract

Neuronal network resilience, the ability of brain circuits to maintain and recover functional connectivity following perturbation, is fundamental to cognitive stability and adaptability. Using human cerebral organoids and multi-electrode arrays (MEAs), we investigated how mechanical stress disrupts network stability and identified key mechanisms regulating recovery. Blast overpressure exposure destabilized small-world network (SWN) organization, increasing network fragmentation and reducing overall integration. Merged SWNs, which exhibit high connectivity, were particularly vulnerable, while fragmented and single SWNs persisted for extended periods, indicating a shift toward less resilient network states. Optogenetic stimulation promoted network recovery, reducing the persistence of fragile states and facilitating transitions toward more cohesive network structures. GABAergic signaling emerged as a critical regulator of network resilience, with pharmacological inhibition exacerbating fragmentation and impairing network reorganization. These findings reveal fundamental principles of how inhibitory networks regulate circuit stability, with implications extending beyond mechanical injury to broader conditions characterized by network dysfunction, including anxiety, depression, PTSD, and neurodegenerative disorders. Understanding the mechanisms governing network adaptation and resilience could inform new therapeutic strategies aimed at stabilizing disrupted neural circuits across a range of neurological conditions.

**Significance statement:** Neuronal networks must dynamically adapt to maintain function in the face of disruption, yet the mechanisms that govern network resilience remain poorly understood. Using human cerebral organoids, we demonstrate that primary blast overpressure destabilizes small-world networks, increasing fragmentation and reducing overall connectivity. Critically, GABAergic signaling emerges as a key stabilizer, with inhibition of GABA receptors amplifying network fragmentation and impairing recovery. These findings provide fundamental insight into how neural circuits resist and recover from mechanical stress, bridging gaps between basic neuroscience, injury pathology, and potential therapeutic interventions. By identifying inhibitory signaling as a regulator of network resilience, our work informs not only traumatic brain injury treatment strategies but also broader efforts to restore functional connectivity in neurological disorders, from epilepsy to neurodegeneration.

## Introduction

The brain operates as a complex, dynamic network, where neurons form circuits that balance local processing with global integration. Small-world networks (SWNs), characterized by high local clustering and short global path lengths, optimize communication efficiency and support cognitive function (Watts and Strogatz, 1998; Bassett and Sporns, 2017). Maintaining the stability of these networks is essential for brain resilience, as network fragmentation or excessive integration can impair information processing and disrupt normal function. Understanding how neural networks respond to perturbations, both mechanical and neurophysiological, is critical for uncovering mechanisms that preserve or restore network stability following disruption.

Neural network resilience can be investigated using controlled experimental models that allow for precise manipulation of network states. Human cerebral organoids, derived from induced pluripotent stem cells (iPSCs), provide a unique platform for studying neural circuit development and adaptation to external stressors (Lancaster and Knoblich, 2014; Birey et al., 2017). These self-organizing three-dimensional cultures recapitulate key aspects of human cortical architecture, including excitatory-inhibitory balance, synaptic activity, and emergent oscillatory dynamics (Trujillo et al., 2019). By leveraging cerebral organoids and multi-electrode arrays (MEAs), we can analyze the principles governing network stability and how external forces affect connectivity.

To probe network resilience, we use two distinct methods that challenge small-world network stability in different ways. Blast overpressure introduces a mechanical perturbation, generating rapid, high-frequency pressure waves that fragment SWNs and reduce overall connectivity (Silvosa et al., 2022). This disruption occurs without immediate cellular damage, making it a valuable model for studying how mechanical stress influences functional network organization (Svetlov et al., 2010; Mac Donald et al., 2011; Rubovitch et al., 2011; Goldstein et al., 2012). Optogenetic stimulation, in contrast, provides a precise way to manipulate neuronal activity and test whether targeted network activation can promote recovery. Previous studies have used optogenetics to map large-scale network effects (Chen et al., 2020) revealing that stimulation can induce both local and brain-wide changes in connectivity. Optogenetic perturbations have also been employed to investigate inhibitory-stabilized networks (ISNs), demonstrating that hippocampal CA1 and CA3 circuits exhibit unexpected stabilization mechanisms, where inhibition paradoxically increases in response to suppressive stimuli (Watkins de Jong et al., 2023). This inhibition-stabilized framework is crucial for network resilience, as perturbations that selectively disrupt inhibitory populations can reveal underlying mechanisms of circuit balance (Sadeh et al., 2017).

In this study, we examine how small-world network stability is shaped by both mechanical and neurophysiological perturbations. Using cerebral organoids as an experimental model, we demonstrate that blast overpressure drives a shift toward network fragmentation and diminished integration, disrupting the intrinsic balance of small-world connectivity. In contrast, optogenetic stimulation counteracts this destabilization, facilitating the recovery of coherent network dynamics and promoting transitions toward more functionally integrated states. Notably, we identify local GABAergic interneuron networks as key regulators of network resilience, dynamically modulating inhibitory-excitatory balance to maintain network stability. These interneuron circuits appear particularly vulnerable to blast overpressure, where their selective disruption precipitates a breakdown in coordinated neuronal activity and amplifies network destabilization. Pharmacological inhibition of GABAergic signaling further exacerbates this fragmentation, reinforcing the critical role of inhibitory circuits in preserving network integrity. Together, these findings provide mechanistic insights into how network-level homeostasis is perturbed by mechanical stressors and suggest potential avenues for targeted interventions aimed at restoring functional connectivity following network disruption.

## Materials and Methods

### Pluripotent stem cell cultures

Human pluripotent stem cells (10121c16; UTSA Stem Cell Core) were cultured under feeder-free conditions on Vitronectin XF™-coated plates (StemCell Technology, #100-0763). The cells were maintained in mTeSR-1 medium (Stem Cell Technologies, 05850) supplemented with 1% penicillin-streptomycin (Life Technologies, 15070).

### Cerebral organoid cultures

Cerebral organoids were generated using previously established protocols. Briefly, pallial and subpallial spheroids were derived from pluripotent stem (PS) cells and assembled in vitro to model the development of the human cerebral cortex.

#### Pallial Spheroid Formation

Dissociated PS cells were plated into ultra-low attachment 96-well plates to form embryoid bodies in neuro induction medium (NIM) containing 20 μM ROCK inhibitor (Y-27632, Stem Cell Technology). The NIM formulation included DMEM-F12 (Invitrogen) with 20% KnockOut Serum Replacement (Invitrogen), GlutaMAX (1:100, Invitrogen), MEM-NEAA (1:100, Gibco), 0.1 mM 2-mercaptoethanol (Gibco), and 1% penicillin-streptomycin (Sigma). For the first six days, neural induction was promoted with 10 μM dorsomorphin (StemCell Technology, 72102) and 10 μM SB-431542 (Tocris). The spheroids were then transferred to neural medium (NM), consisting of Neurobasal A (Gibco, 10888) with B27 supplement (−Vitamin A, Invitrogen), GlutaMAX (1:100, Invitrogen), and 1% penicillin-streptomycin (Sigma). To support neural progenitor expansion, NM was supplemented daily with 20 ng/mL fibroblast growth factor (Peprotech) and 20 ng/mL epidermal growth factor (Peprotech) for 10 days, then every other day for 9 days. For neural differentiation, NM was supplemented every other day with 20 ng/mL brain-derived neurotrophic factor (Peprotech) and 20 ng/mL NT3 (StemCell Tech) until Day 43. After Day 43, spheroids were maintained in NM without additional factors.

#### Subpallial Spheroid Formation

Subpallial spheroids were prepared similarly to pallial spheroids but with pathway-specific modifications. From Day 4, Wnt pathway inhibition was induced by adding 5 μM IWP-2 (StemCell TEch) to NIM and NM. From Day 12 to Day 24, sonic hedgehog pathway activation was achieved using 100 nM SAG (StemCell Tech) in combination with IWP-2. After Day 24, subpallial spheroids were maintained under the same conditions as pallial spheroids.

#### Cerebral Organoid Fusion

To replicate human cortical development, pallial and subpallial spheroids were fused. On Day 43, spheroids were placed together in a single well of a 24-well plate and incubated at an angle for 5-7 days, with media changes every 4 days. Once fusion was complete, the organoids were transferred to a rotating shaker in the incubator, and NM without additional factors was replaced three times per week.

### Blast overpressure exposure

A previously developed tabletop “blast” device was used to deliver quasi-hydrostatic pressure waves to cerebral organoids (Vidhate et al., 2021; Silvosa et al., 2022). Pressure loading in this setup is considered quasi-hydrostatic because the pressure waves travel at the speed of sound in water (∼1480 m/sec), and the pressure chamber is approximately 30 mm in length. This setup ensures that any pressure changes homogenize within ∼20 µs, corresponding to a characteristic frequency of ∼50 kHz, which is 10 times faster than the highest frequency used in the experiment (5 kHz). To expose the organoids to high-frequency pressure waves, pre-warmed, CO2-buffered neurobasal medium (3-5 mL) was added to the blast chamber. Organoids from each experimental group were placed together in the chamber, which was sealed to avoid air bubbles, then installed into the support frame for pressure wave application.

### Spatial distribution analysis

To assess the spatial organization of neuronal subtypes in cerebral organoids, we utilized custom Python scripts to calculate Ripley’s *K*-function, the Nearest Neighbor Distribution (*G*-function), and the transformed Ripley’s *K*-function (*L*-function). These functions were calculated from regions of interest (ROIs) measured using ImageJ (NIH), which provided the spatial coordinates of labeled cells for each neuronal subtype.

#### Ripley’s K-Function

Ripley’s *K*-function evaluates the extent of spatial clustering or dispersion by comparing the observed distribution of cells to a theoretical random distribution (CSR: Complete Spatial Randomness). For each radius (*d*), the number of neighboring cells within *d* of a given cell was calculated, normalized by the overall density of cells in the ROI. The equation used is:

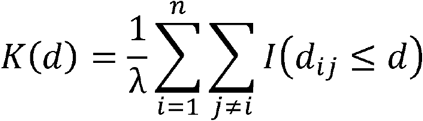

where λ is the overall density of cells, *d*_ij_ is the pairwise distance between cells *i* and *j*, and *I(d*_ij_ *< d)* is an indicator function that equals 1 if the distance between *i* and *j* is less than or equal to *d*, and 0 otherwise. The CSR line *(πd*^2^) was used as a reference to identify clustering (*K(d) > rrd*^2^) or dispersion (*K(d) < πd*^2^).

#### Nearest Neighbor Distribution (G-Function)

The *G*-function, or Nearest Neighbor Distribution, measures the probability of finding at least one neighboring cell within a given radius (*d*). It is calculated as:

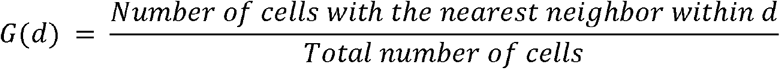

The *G*-function was used to identify the density of cells at small spatial scales. A steeper slope of *G(d)* indicates higher local density, while a more gradual slope reflects greater dispersion.

The radius at which *G(d) = 1* was recorded as the distance at which all cells had at least one neighboring cell.

#### Transformed Ripley’s K-Function (L-Function)

The *L*-function normalizes Ripley’s *K*-function to facilitate the interpretation of deviations from CSR. The transformed equation is:

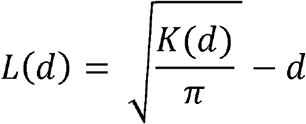

Values of *L(d) > 0 i*ndicate clustering, and values of *L(d) < 0* indicate dispersion. The *L*-function was used to quantify the maximum deviation from CSR and calculate the clustering area (cumulative positive deviations of *L(d)* over the range of radii).

For each neuronal subtype (TUJ1, SST, PV, and CTIP2), the measured ROIs were analyzed using these functions, and metrics such as maximum deviation from CSR, clustering area, and the radius at *G(d) = 1* were extracted. All analyses were performed using custom Python scripts, and results were visualized to compare clustering behavior across neuronal subtypes.

### MEA recording and optogenetic stimulation

Cerebral organoids were plated onto a 24-well MEA plate (24W700/100F-288, Multichannel Systems) pre-coated with Vitronectin XF™ (Stem Cell Technology, #07180). Each well contained 12 gold electrodes, 100 μm in diameter, spaced 700 μm apart. Prior to plating, fresh neurobasal medium (NM) was added to each well and incubated in 5% CO2 for 30 minutes to allow buffering. MEA recordings were performed using the Multiwell-Screen Acquisition software (Multichannel Systems, v1.11.7.0). Organoids were plated immediately after the “blast” procedure, and the plate was placed on the MEA system for 1 hour before the initial recording. Throughout the experiment, temperature was maintained at 37°C with 5% CO2. All recordings were sampled at 10 kHz, with a band-pass filter of 1 Hz-3500 Hz applied.

#### Channelrhodopsin expression and optogenetic stimulation

Low-frequency optogenetic stimulation of neural networks was achieved using Cre-dependent, AAV-mediated viral transduction of Channelrhodopsin. Mature cerebral organoids (180 days old) were infected with AAV1-hSyn-eGFP-Cre (Addgene #105540) at a final titer of 7.5e11 overnight. The following day, fresh media was added and exchanged daily for 7 days to ensure all viral particles were removed. GFP expression was confirmed within 10 days post-infection. To specify expression in neurons, a secondary virus, AAV1-CAG-FLEXFRT-ChR2(H134R)-mCherry (Addgene #75470), was added at a final titer of 1.3e12. Experiments were conducted once both mCherry and eGFP expression were confirmed.

Pulsed blue LED light was delivered using a controlled multiwell LED system (MW24-opto-stim, Multichannel Systems). Every 10 seconds, a 5 ms light pulse was applied over a 60-second period to provide a synchronizing stimulus to the organoids.

#### Synaptic pharmacology

Optogenetic stimulation was paired with pharmacological agents to modulate synaptic transmission. An initial 60 seconds of baseline activity was recorded, followed by the application of AMPA/CNQX (50/10 µM) to block glutamatergic synaptic transmission, and a 2-minute recording with LED activation was obtained. After washout, Gabazine (100 µM) was added to inhibit GABA_A_ receptor function, followed by another 2-minute recording. Lastly, lidocaine (100 µM) was used to broadly block sodium channel activity after the final washout, with a subsequent recording to assess the effects on network activity.

### Data Acquisition and Organization

Neuronal data were collected from multiple experimental wells, each containing electrode arrays to record neuronal activity. Raw data were stored in .h5 file format, with each file corresponding to a specific experimental condition. The data were organized into three directories: a Data directory containing all .h5 files, an Origin Directory for intermediate CSV files, and a Destination directory for the final output files, including synchronization maps, heatmaps, and other analyses. Data preprocessing was initiated by extracting analog streams from the .h5 files using the McsPy package (McsPy, 2023). Each .h5 file was read and segmented based on the experimental condition, and the segmented data were saved as CSV files. These CSV files contained the raw unsegmented data for each recording channel and were further divided into smaller segments based on a predefined segment length to facilitate subsequent analysis.

### Generation of Synchrony maps

#### Calculating Synchronization Index for Network Analysis

Synchronization indices (SI) were computed for each pair of channels to quantify the degree of synchronization between neural signals as previously published (Silvosa et al., 2022). This was achieved using a complex Morlet wavelet transform applied to the time-series data of each channel pair, followed by the calculation of phase synchronization values. The SI was defined as the absolute value of the mean vector of the Euler representation of phase angle differences between the signals. Network synchrony maps were generated by forming matrices from the synchronization indices and clustering coefficients. Nodes were represented as circles in these maps, with their size proportional to the number of connections, and their color indicating the magnitude of the clustering coefficient. Connections between nodes were depicted as lines, with their thickness proportional to the synchronization index.

#### Clustering Coefficient Calculation

Let *C*_*ij*_ represent an existing connection (i.e. Synchronization Index above average) between node *i* and node *j*. The following set, *N*_*i*_, then, expressed all nodes, *n*_*j*_, that are connected to node *i*, where *C* represents a set of all existing connections within the network:

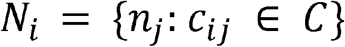

For convenience, |*N*_*i*_| = _*i*_. If node *i* is connected to 3 nodes, *l*_*i*_ = 3. For node *i*, a connection *C*_*ij*_ is referred to as a direct connection, while connection *c*_*jk*_, where both *n*_*j*_ and *n*_*k*_ are elements of *N*_*i*_, is an indirect connection. The clustering coefficient (*CC*_*i*_) of each node *i* within the neuronal network was calculated to quantify the degree of connectivity between the nodes in the network. The clustering coefficient was determined using the formula:

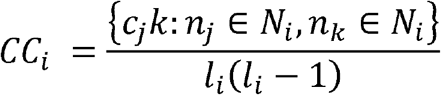

A high *CC* suggests that the nodes connected to node *i* are all strongly connected to each other. For a node with a clustering coefficient of 1, i.e. every possible indirect exists. We call a node “significant” if its *CC* is above average for its phase. A node is “perfect” if its *CC* is equal to 1. By the range of a *CC*, all perfect nodes are significant, but not all significant nodes are necessarily perfect.

#### Independent Network Identification

Independent networks were identified based on the presence of perfect nodes (*CC*=1). For each perfect node, all directly connected nodes were identified, forming a network. If multiple perfect nodes were present within a phase and were interconnected, they were the same independent network. The number of independent networks and perfect nodes were calculated and recorded for each experimental phase.

### Network transition states

Single SWNs were defined by the presence of a single central node with a CC = 1, indicating complete interconnectivity with its neighboring nodes. Additionally, all directly connected nodes were required to have a synchronization index (SI) exceeding a set threshold to qualify as significant connections. Fragmented SWNs consisted of multiple independent SWNs, each with a central node with CC = 1, but the connections between nodes did not meet the SI threshold, indicating limited synchronization. Merged SWNs were identified when multiple independent SWNs, each with CC = 1, had an SI > threshold, resulting in the formation of an integrated, larger network structure. In contrast, fragile networks lacked any nodes with CC = 1, signifying minimal or absent interconnected clusters and representing a less stable network state.

To quantify transitions between network states, data from microelectrode array (MEA) recordings were analyzed phase by phase using a custom Python script. The dataset, containing network state information for each phase and well, was imported from a CSV file. Transitions between consecutive states (e.g., X → S) were identified and counted using a transition-counting function applied sequentially across 1-second phases. This function iterated through the data, recording each current and subsequent state to compile the frequency of each transition type. The aggregated results were organized into a DataFrame, which displayed the transition counts for each well. The DataFrame was then printed for verification and exported to an Excel file for further analysis and reporting.

### Experimental Design and Statistical Analysis

All data are presented as mean□±□standard error of the mean (SEM) unless otherwise specified. Statistical comparisons were performed using two-tailed Student’s t-tests or analysis of variance (ANOVA) for datasets with equal variances. Post hoc analyses following ANOVA were conducted using Tukey’s multiple comparisons test to identify differences between groups, indicated by an asterisk and black bar. Data collection and analysis were conducted simultaneously to ensure consistency. Statistical analyses were performed using GraphPad Prism (v.8.4.3) in accordance with the GraphPad Prism Statistics Guide.

### Data and Code Availability

All data supporting the findings of this study are included within the paper. Additional information is available upon reasonable request to the corresponding authors. Custom MEA analysis code was written in Python (v.3.10.1) and is available upon request (Lybrand, 2025).

## Results

### Neurons distributed in cerebral organoids exhibit a hierarchical organization to network structure

Cerebral organoids, three-dimensional cultures that replicate brain development, form complex network functions similar to those found in neonatal brain waves (Trujillo et al., 2019). To explore how complex neural circuits in these organoids respond to mild primary blast overpressures, we utilized human induced pluripotent stem cells, maturing them into cerebral organoids over six months using a guided, dual SMAD inhibition protocol (Birey et al., 2017). These organoids exhibited expanse neuronal networks observed by the express of beta-III tubulin (TUJ1), a neuronal specific microtubule protein. Additionally, key cell populations from a cortical niche are developed, including deep layer cortical neurons marked by COUP-TF-interacting protein 2 (CTIP2), parvalbumin (PV) and somatostatin (SST) subtype interneurons. (Figure 1 A-F). These excitatory and inhibitory neuronal subtypes occurred in similar frequency across organoids in the study, however, their distribution throughout the organoid varied.

**Figure 1.**
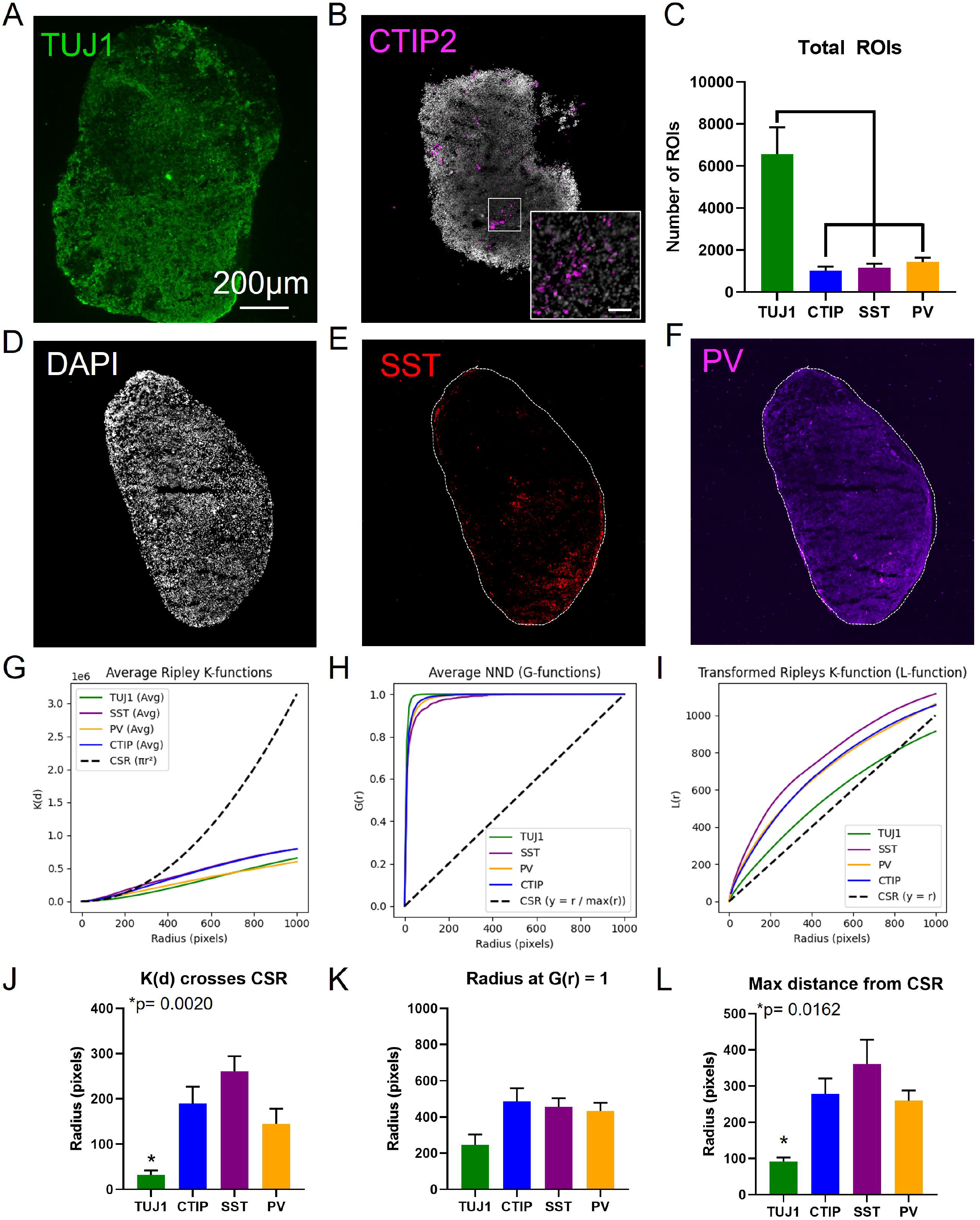
Structural organization of neuronal networks within cerebral organoids. (A) Representative image showing the expression of beta-III tubulin (TUJ1) to visualize the global distribution of neurons within cerebral organoids. (B) Representative image of CTIP2 expression marking cortical neuron subtypes. Inset shows a magnified view (scale bar = 50 μm). (C) Quantification of the total number of ROIs identified for analysis across neuronal subtypes. (D-F) Representative images of nuclei and interneuron subtypes in a single organoid: (D) DAPI staining for nuclei distribution, (E) somatostatin (SST)-positive interneurons, and (F) parvalbumin (PV)-positive interneurons. (G) Quantification of spatial clustering using Ripley’s K-function. A K(d) value above the CSR line indicates clustering, while a value below suggests dispersion. (H) Nearest Neighbor Distribution (NND) or G-function quantifying the density of neighboring cells at increasing radii. (I) Transformed Ripley’s K-function (L-function), highlighting deviations from CSR to assess clustering intensity. (J) Quantification of the radius at which the K(d) values in (G) cross the CSR line, highlighting spatial organization differences. (K) Quantification of the radius at which G(r) reaches 1 across neuronal subtypes, reflecting the distance at which every neuron has at least one neighbor. (L) Quantification of the maximum distance L(r) deviates from the CSR line, representing the strength of local clustering. All data are presented as mean ± SEM, with n = 12 for each group. Exact p-values from ANOVA are shown in (J) and (L), with asterisks (*) indicating statistically significant differences from multiple comparisons.

To analyze the structural organization of the neuronal networks, spatial distribution was assessed using Ripley’s K-function, which suggested differences in clustering behavior across neuronal subtypes. A K(d) value above the CSR line indicates clustering, while a value below suggests dispersion or regular spacing. TUJ1-positive neurons, representing the general neuronal population, displayed the weakest clustering, with K(d) values consistently positioned below the CSR (Complete Spatial Randomness) line (Figure 1G). This implies a more uniform distribution, where neurons are more evenly spaced than randomness would predict, likely due to repulsive interactions or mechanisms that maintain distance between neurons to optimize network functionality and prevent overcrowding. In contrast, CTIP2, SST, and PV neurons demonstrated stronger clustering, with K(d) values exceeding the CSR line over a range of radii. Among these subtypes, SST neurons exhibited the strongest clustering behavior, followed closely by CTIP2 neurons, while PV neurons showed intermediate clustering. These differences in clustering behavior were further quantified using the maximum distance from the CSR line and the clustering area to quantify clustering (Figure 1J). SST neurons displayed the highest maximum distance and clustering area, indicative of their robust local clustering, while CTIP2 and PV neurons followed with progressively lower values. TUJ1-positive neurons showed the smallest clustering area and maximum distance, consistent with their more uniform distribution.

The Nearest Neighbor Distribution (NND) or G-function provided additional insights into clustering density (Figure 1H). The slope of G(r) reflects the rate at which neurons accumulate neighbors as the radius increases: a steeper rise in G(r) indicates higher density and shorter nearest-neighbor distances, while a gradual rise reflects greater dispersion. TUJ1 neurons exhibited the steepest slope, suggesting a rapid accumulation of neighbors at small radii, indicative of a densely packed network. In contrast, SST neurons showed the most gradual slope, consistent with a more dispersed spatial arrangement and slower accumulation of nearest neighbors. CTIP and PV neurons displayed intermediate slopes, reflecting a balance between localized clustering and broader spacing.

Quantification of the radius at which G(d)=1 (i.e., the point where every neuron has at least one neighbor) provided additional insight into the extent of spatial dispersion (Figure 1K). Despite their steep slope in Figure 1H, TUJ1 neurons reached G(d)=1 at the smallest radius (∼200 pixels), confirming their dense clustering and close nearest-neighbor distances. In contrast, SST, CTIP, and PV neurons required significantly larger radii (∼500 pixels) to achieve G(d)=1, reflecting greater overall dispersion within the organoid. Notably, the similarity in radii for CTIP and PV neurons suggests comparable spatial organization, while SST neurons, despite their gradual slope in Figure 1H, reached G(d)=1 at a radius similar to CTIP and PV neurons, highlighting their broader but still localized distribution.

Further validation came from the transformed Ripley’s K-function (L-function), which normalizes the K-function to highlight deviations from CSR (Figure 1I). Values above the CSR line indicate clustering, with greater deviations signifying stronger clustering. SST neurons displayed the largest deviations from CSR, indicating robust local clustering, followed by CTIP2 neurons. PV neurons exhibited moderate clustering, while TUJ1-positive neurons showed the weakest clustering, with values remaining near the CSR line. Quantitative metrics supported these observations, with SST neurons showing the largest maximum distance from CSR and clustering area, indicative of strong local clustering. CTIP2 neurons also displayed substantial clustering, while PV neurons exhibited intermediate clustering behavior, and TUJ1-positive neurons showed the lowest clustering metrics (Figure 1J-L).

Together, these analyses highlight a hierarchical spatial organization of neuronal subtypes within cerebral organoids, revealing distinct clustering behaviors that likely reflect their functional roles in network architecture. TUJ1-positive neurons, representing the general neuronal population, exhibited the weakest clustering and most uniform distribution, as evidenced by their consistently low deviations from CSR and the smallest clustering area. This arrangement facilitates broad connectivity across the organoid. In contrast, SST neurons demonstrated the strongest clustering, with the highest maximum distance from CSR, the largest clustering area, and a gradual rise in the NDD function, reflecting their role in forming dense, localized inhibitory networks. CTIP2 and PV neurons displayed intermediate clustering, balancing local connectivity with broader spatial organization. These findings demonstrate that cerebral organoids replicate the spatial complexity of cortical niches, balancing local clustering and global dispersion to support network functionality. This insight provides a foundation for understanding how structural organization underpins network behaviors and evaluating the impact of experimental perturbations like mild blast overpressure.

### Emergent behaviors of neuronal networks developed in cerebral organoids

To assess the impact of blast overpressure on network synchrony dynamics, cerebral organoids were exposed to a high-frequency overpressure wave (250 kPa at 5000 Hz), designed to disrupt neuronal activity without causing immediate cellular damage (Silvosa et al., 2022). Control organoids, which were not exposed to the blast, and blast-exposed organoids were immediately plated onto a multielectrode array (MEA). Neural activity was recorded 1-hour post-blast (Supplemental Figure 1). We analyzed 2 minutes of raw MEA data and applied Morlet wavelet convolution to extract time-frequency information across a range of frequencies. The data were segmented into 1-second epochs, resulting in 120 epochs, and the synchrony index was calculated for each epoch to assess the degree of synchronization between channel pairs.

We developed network synchrony maps to visually represent the dynamics of synchronization within the network (Figure 2A). These maps illustrate the strength of phase synchronization between channel pairs, with thicker lines indicating stronger synchronization. Nodes with more connections, reflected by the size of the circles, were more extensively involved in network-wide synchronization. In parallel, clustering coefficient (CC) analysis quantified local connectivity within the network. Nodes with higher CC values were more likely to form tightly interconnected clusters, indicative of localized synchrony. Notably, nodes with a CC of 1 demonstrated complete interconnectivity with their neighbors, reflecting the presence of highly synchronized subnetworks across phases. To improve clarity in understanding prominent networks, synchrony network maps were reduced to significant connections only to produce a simplified network map (Supplemental Figure 2). These simplified network maps were used for all data analysis and display only include the nodes with a CC = 1, and all associated connections (Supplemental Video 1).

**Figure 2.**
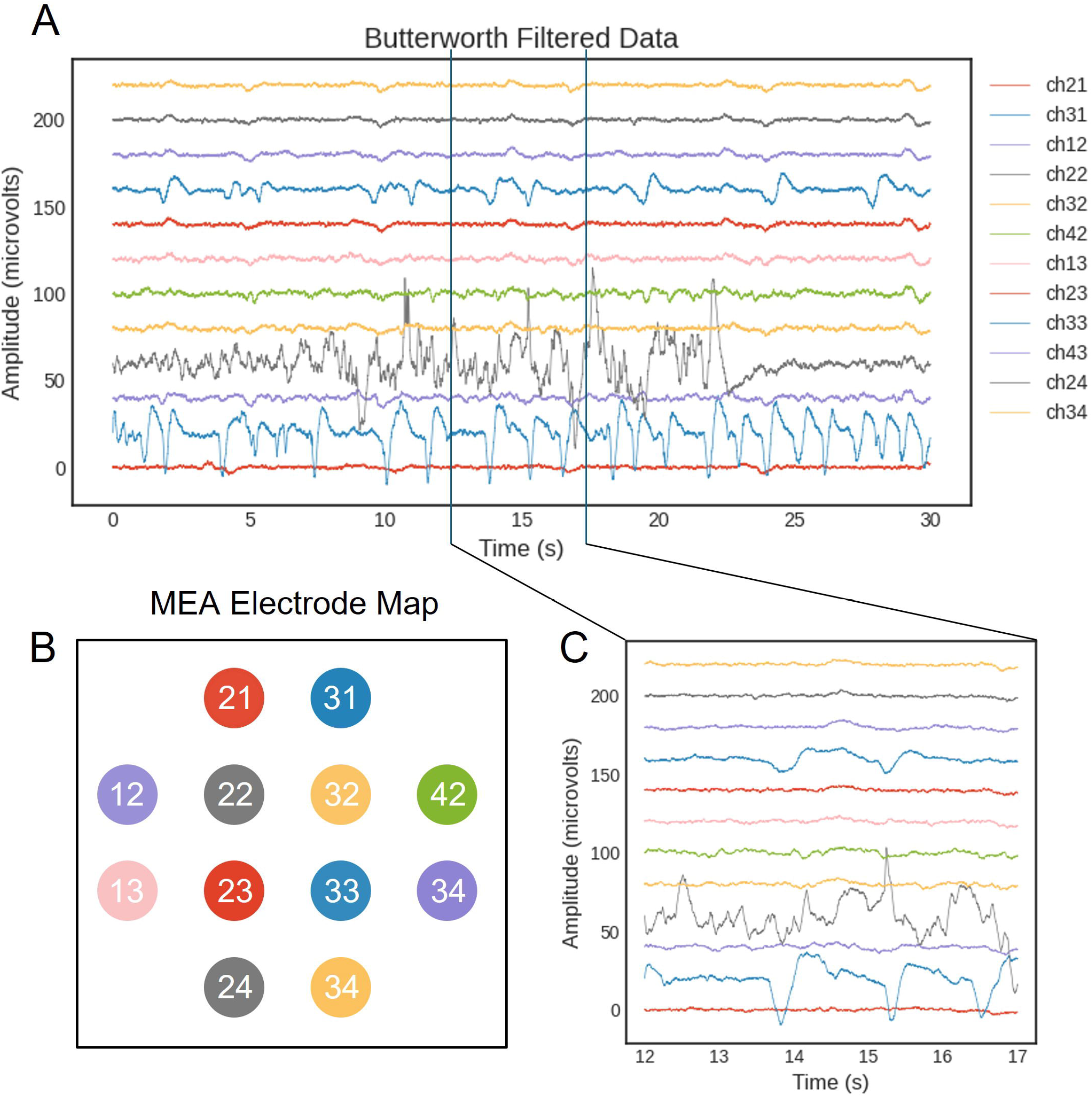
Spatial networks of MEA. (A) Representative raw data from multielectrode array with an applied Butterworth filter. With an increased timescale found in in panel C. (B) A color coded electrode map shows the spatial orientation of all 12 channels that correspond to the channels displayed in panel A.

Within our MEA data, we identified four distinct small-world network (SWN) states. SWNs are networks that combine high local clustering with short average path lengths, allowing for efficient information transfer within a network. They are particularly important because they optimize both local processing and global communication. Single SWNs (green) were characterized by a single central node with several directly connected nodes that exhibited strong synchronization. Fragmented SNWs (orange) consisted of independent SWNs, each with a central node, but lacking significant synchronization between the central nodes. Merged SWNs (blue) were identified when multiple SNWs became significantly synchronized, indicating the merging or expansion of several networks into a larger, more integrated structure. Lastly, fragile networks (black) were defined by the absence of any nodes with a clustering coefficient (CC) of 1, indicating a lack of highly interconnected nodes (Figure 2B).

Out of the total 120 epochs, we quantified an average of 43.5 ± 2.3 single SWNs from control organoids and 42.0 ± 1.8 from blast organoids (Figure 2C) that were not significantly different (p=0.6269, unpaired t-test, n=4 each). We observed an increase in the number of fragmented SWNs following blast (Figure 2D, 12.0 ± 2.0 control and 21 ± 1.6 blast; p=0.0115, unpaired t-test), and a significant decrease in the number of merged SWNs (Figure 2E, 19.75 ± 2.4 control and 13.0 ± 1.2 blast; p=0.0459, unpaired t-test). There were no observed differences in the frequency of fragile networks following blast (Figure 2F, 45.25 ± 3.2 control and 37.0 ± 2.5 blast; p=0.5234, unpaired t-test). Altogether, neuronal networks in cerebral organoids spend most of the time in a SWN or fragile network state, and these states appear largely unaffected by blast overpressures. In contrast, fragmented and merged SWNs are selectively affected by blast overpressures, with an increase in fragmentation and a decrease in merged SWNs, suggesting network disintegration and a loss of connectivity.

For each network state, we measured the number of connections for each significant node. The frequency distributions of node connections within SWNs were analyzed and visualized under both Control and Blast conditions (Figure 2G). The plot reveals the number of nodes within each network type that exhibit specific numbers of connections, ranging from 0 to 8.

Under the control scenario, the single SWNs displayed a peak at 2 connections and gradually declined in frequency as the number of connections increased, indicating a moderate level of connectivity typical of a sparsely connected network. The fragmented SWN, similarly, showed a peak at 2 connections but maintained a slightly higher frequency of nodes with 3 and 4 connections compared to the single SWN, suggesting a slightly more connected network structure. The merged SWN, on the other hand, exhibited a broader distribution with the highest frequencies occurring between 2 and 5 connections, reflecting a more densely connected network topology.

Following exposure to a blast, all network types demonstrated significant changes in their connectivity patterns. Single SWN under blast conditions showed an increased frequency of nodes with 2 connections and a notable presence of nodes with 3 connections compared to the control. This suggests a reorganization towards slightly higher connectivity post-blast.

Fragmented SWN displayed a remarkable increase in nodes with 2 to 4 connections, highlighting a shift towards greater connectivity and possibly indicating adaptive changes in network structure to maintain functionality. The merged SWN showed the most pronounced changes with a more uniform distribution across 2 to 6 connections, suggesting extensive reorganization and increased redundancy within the network.

Furthermore, blast overpressure did not significantly affect the average number of connections in single SWNs (2.9 ± 0.07 in control vs. 2.7 ± 0.13 in blast conditions) or fragmented SWNs (2.9 ± 0.08 in control vs. 2.98 ± 0.07 in blast conditions). However, blast exposure led to a significant reduction in the number of connections in merged SWNs (4.23 ± 0.13 in control vs. 3.69 ± 0.09 in blast conditions, *p < 0.0001, F(5, 1058) = 17.98, One-way ANOVA with multiple comparisons).

Altogether, blast overpressure appears to selectively affect the more integrated, highly connected merged SWNs, leading to a significant reduction in their connectivity. In contrast, single and fragmented SWNs seem to be more resilient, showing no significant changes in their average number of connections. This suggests that the blast primarily disrupts the most connected and integrated networks, which may contribute to the overall loss of network integration observed after trauma.

### Optogenetic activation stabilizes network transition states

To assess network stability, we applied optogenetic stimulation to cerebral organoids at a controlled frequency to measure dynamic changes in network states over time (Figure 3). Network stability was defined by the network’s ability to maintain or transition into organized states that support efficient local and global communication. Network states were categorized for each 1-second phase and visualized using a heatmap. These states included Fragile (X), representing disorganized and low-connectivity configurations; Fragmented (F), characterized by independent clusters with limited synchronization; Single (S) small-world networks (SWNs), defined by a central node with strong, connected interactions; and Merged (M) SWNs, indicative of broader integration and highly synchronized clusters (Figure 3A). A transition matrix was generated to quantify how frequently each network state transitioned into another in the subsequent phase (Figure 3B). At baseline, X→X transitions were the most frequent, indicating that the networks tend to return to a fragile state. Additionally, frequent transitions between X and S states were observed, suggesting that the network often fluctuates between unstable and more stable configurations.

**Figure 3.**
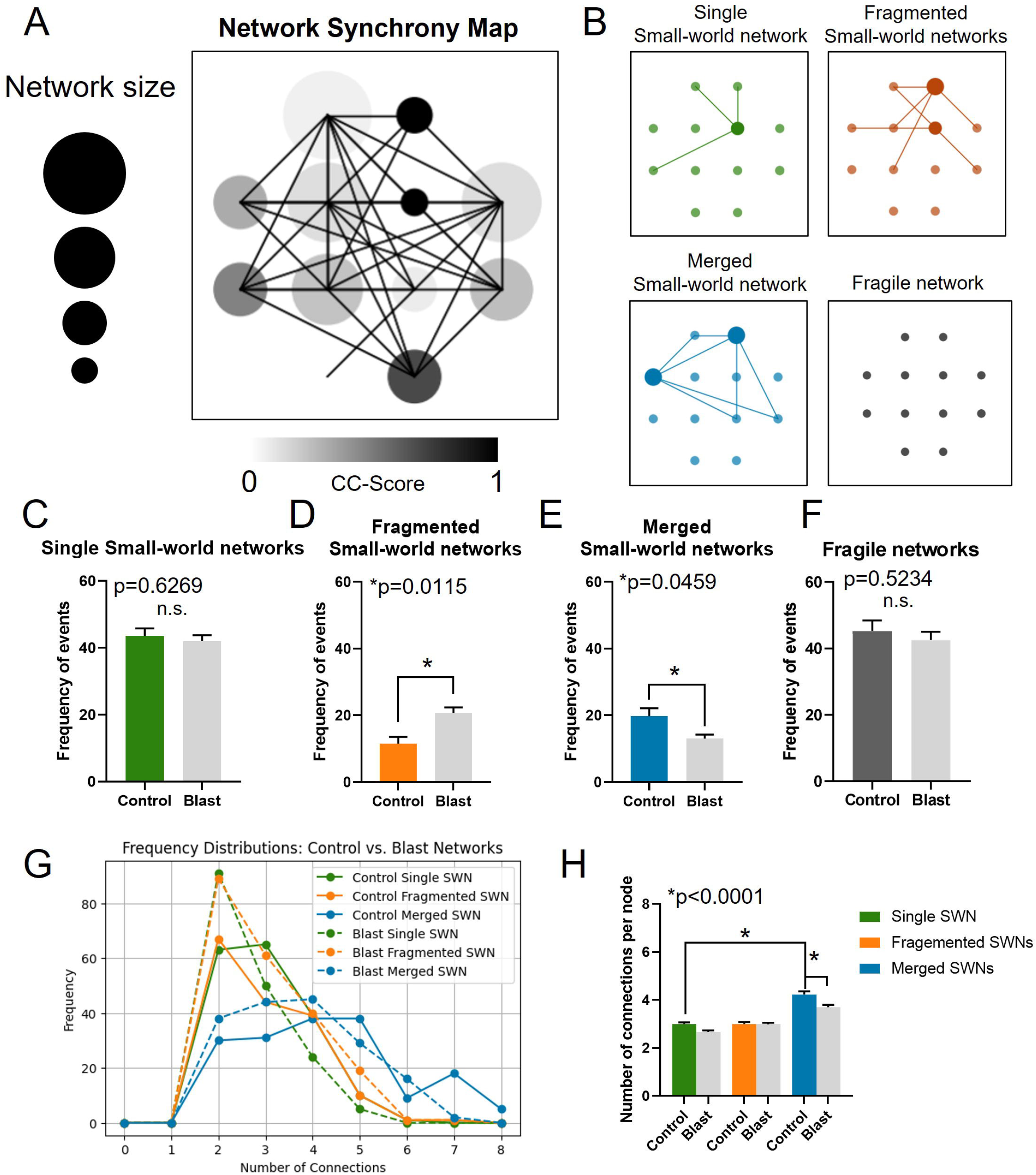
Synchrony maps reveal emergent network properties in cerebral brain organoids. (A) Representative network synchrony map illustrating the emergent properties of neuronal networks. Node size reflects the number of network connections, with larger nodes having more connections. Node shading corresponds to the clustering coefficient (CC), with darker shading indicating stronger connectivity. Thicker lines between nodes represent stronger connections. (B) Simplified representations of significant nodes and their connections depict four distinct network states: Single Small-World Networks (SWNs, green), Fragmented SWNs (orange), Merged SWNs (blue), and Fragile networks (black). (C-F) Quantification of the frequency of each network state during the recording period for control and blast conditions: (C) Single SWNs, (D) Fragmented SWNs, (E) Merged SWNs, and (F) Fragile networks. (G) Frequency distributions comparing the change in the number of connections across network states between control (solid lines) and blast (dashed lines) conditions. (H) Quantification of the number of connected nodes across each network state, comparing control and blast conditions. All data are presented as mean ± SEM, with n = 12 per group. Exact p-values from t-tests (C-F) and ANOVA with multiple comparisons (H) are reported, with asterisks (*) denoting statistically significant differences.

Low-frequency stimulation of the networks was achieved through AAV-mediated viral transduction of Channelrhodopsin under the synapsin promoter. Pulsed blue LED light was delivered every 10 seconds to provide a synchronizing stimulus (Figure 3C and D). Network state transitions were quantified at both baseline and during light stimulation.

Following light stimulation, we observed a significant increase in F→S transitions (from 2.75 to 5.5, *p* = 0.0488, F(15, 96) = 17.72, two-way ANOVA with multiple comparisons; Figure 3E, G) and a decrease in X→X transitions (from 10.75 to 5.25, *p* = 0.0028, F(15, 96) = 17.72, two-way ANOVA with multiple comparisons; Figure 3E, F). The increase in F→S transitions indicate enhanced network synchronization, shifting from fragmented to single small-world networks (SWN), suggesting improved network stability. Meanwhile, the decrease in X→X transitions reflect a reduction in the persistence of the fragile state, further indicating improved resilience and network stabilization.

### Blast overpressure destabilizes network transition states

In a parallel experiment, cerebral organoids were exposed to blast overpressure, and transition state dynamics were mapped to visualize changes in network state transitions (Figure 4A). At baseline post-blast, we predominantly observed transitions from S→X (Figure 4B). Although there was a modest increase in S→X transitions between control and blast at baseline, this increase was not statistically significant (Supplemental Figure 3). Following blast exposure, only two transition states were significantly affected. There was a decrease in X→X transitions (from 10.75 to 7.0, p = 0.0249, F(15, 96) = 24.50, two-way ANOVA with multiple comparisons), which remained the most frequent transition across both datasets (Figure 4E, F). Additionally, there was a significant decrease in F→M transitions (from 2.00 to 0.50, p = 0.0496, F(15, 96) = 24.50, two-way ANOVA with multiple comparisons), though this transition was one of the least frequent overall. Overall, the data suggest that blast overpressure impairs the network’s ability to stabilize and synchronize, leading to increased transitions between unstable states and a decreased capacity to reorganize into a more synchronized, coherent network. The decrease in F→ M transitions further highlights the loss of functional recovery mechanisms, while the reduction in X→X transitions suggests increased network dynamics without meaningful stability.

**Figure 4.**
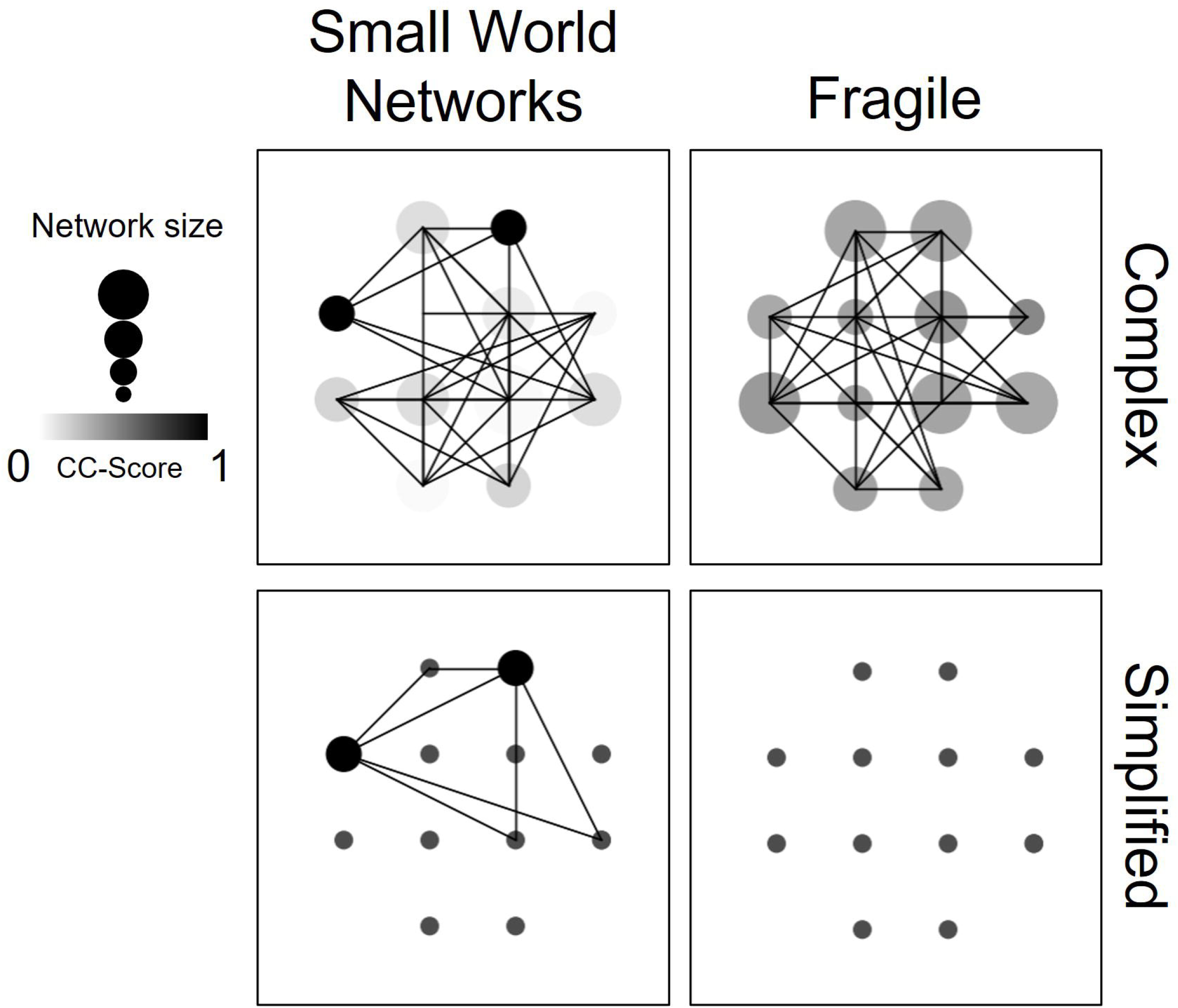
Defining robust and fragile networks. Visual representation of a small world network map with all available connections (top left) and fragile network (top right) with no significant nodes present. Bottom panel represents simplified network map that removes all non-significant connections. On bottom left, only significant nodes and connections remain. On the bottom right, in fragile networks, no significant nodes are present.

To assess how network stability responds to stimulation, pulsed light was applied at a low frequency to activate the network (Figure 4C). Similar to the no-blast control, we observed a decrease in X→ X transitions following stimulation (Figure 4D). The most significant change across all transition states was a robust increase in S→ S transitions (from 6.00 to 10.50, p = 0.0128, F(15, 96) = 17.21, two-way ANOVA with multiple comparisons) (Figure 4E). Altogether, pulsed stimulation significantly enhances the stability and resilience of networks exposed to blast overpressure. The decrease in X→ X transitions and the increase in S→S transitions indicate that stimulation helps the network escape fragility and maintain synchronization, offering insights into potential mechanisms of network recovery post-injury.

### GABAergic regulation of network stability and fragmentation

Acute changes in network stability may result from rapid shifts in synaptic function. To investigate the synaptic mechanisms underlying network transition states, mature cerebral organoids were exposed either to blast overpressure or served as no-blast controls. Organoids were plated for MEA recordings, and optogenetic stimulation was applied to promote network stability and assess baseline function.

To determine whether synaptic activity contributes to network fragmentation, we introduced pharmacological agents targeting different aspects of synaptic function: AMPA/CNQX (50/10 µM) to block glutamatergic synaptic transmission, Gabazine (100 µM) to inhibit GABA_A_ receptor function, and lidocaine (100 µM) to block sodium channel activity more broadly (Figure 5A). After recording a 2-minute baseline response with light stimulation, a wash-in period for each drug was conducted. Following the wash-in, a 2-minute stimulated response was recorded in the presence of each agent. Washout periods between drug applications ensured network recovery before the subsequent agent was administered.

**Figure 5.**
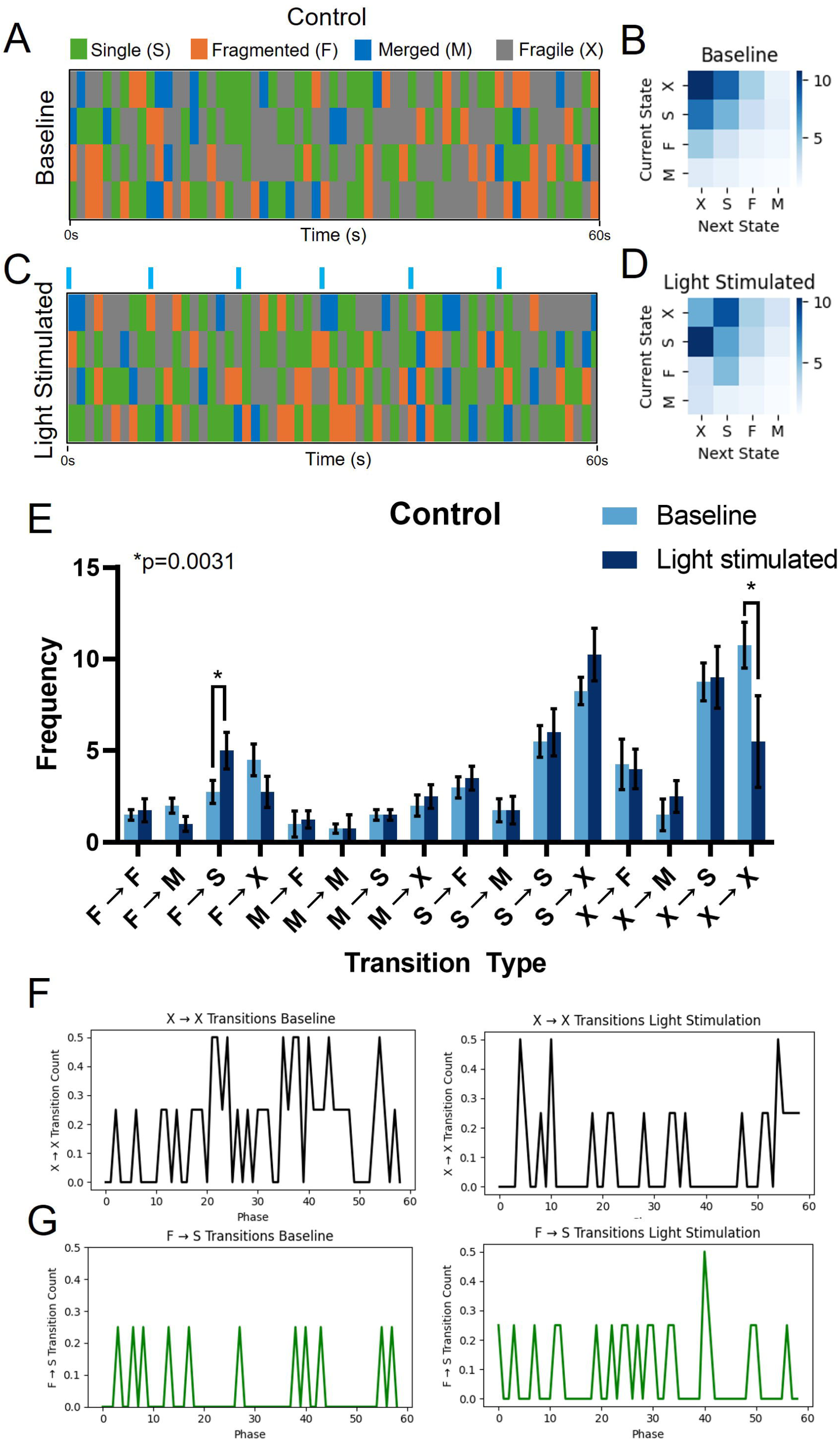
Optogenetic activation stabilizes network transition states. (A) Visualization of network state transitions at baseline. Single SWNs are represented in green, fragmented in orange, merged in blue, and fragile states in grey. Each row represents an individual sample. (B) Quantification of network state transitions at baseline. (C) Visualization of network state transition in response to light stimulation using optogenetics. Blue bars above the plot indicate the time point blue light was pulsed to coordinate network activation. (D) Quantification of network state transitions with light stimulation. (E) Histogram to quantify the number of each transition types between baseline (light blue) and light stimulated (dark blue). (F) Individual representative example of X→X transitions between baseline, on the left, and the response to light stimulation, right. (G) Individual representative example of F→S transitions between baseline, on the left, and the response to light stimulation, right. All data are presented as mean ± SEM, with n = 4 per group. Exact p-values from ANOVA with multiple comparisons are reported, with asterisks (*) denoting statistically significant differences.

**Figure 6.**
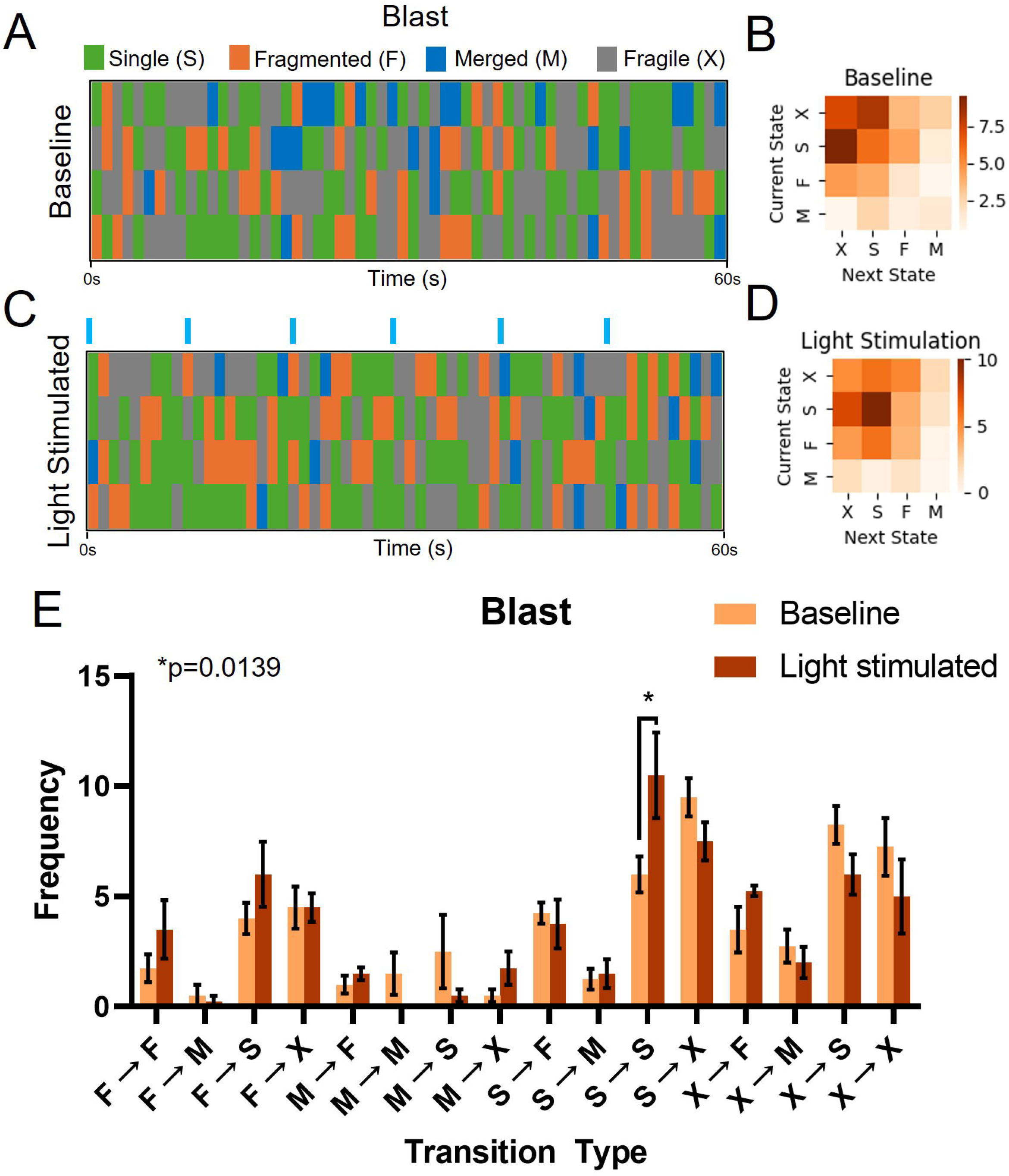
Blast overpressure destabilizes network transition states. (A) Visualization of network state transitions at baseline following blast. Single SWNs are represented in green, fragmented in orange, merged in blue, and fragile states in grey. Each row represents an individual sample. (B) Quantification of network state transitions at baseline following blast. (C) Visualization of network state transition in response to light stimulation using optogenetics following blast. Blue bars above the plot indicate the time point blue light was pulsed to coordinate network activation. (D) Quantification of network state transitions with light stimulation following blast. (E) Histogram to quantify the number of transition types between baseline (light orange) and light stimulated (dark orange). All data are presented as mean ± SEM, with n = 4 per group. Exact p-values from ANOVA with multiple comparisons are reported, with asterisks (*) denoting statistically significant differences.

**Figure 7.**
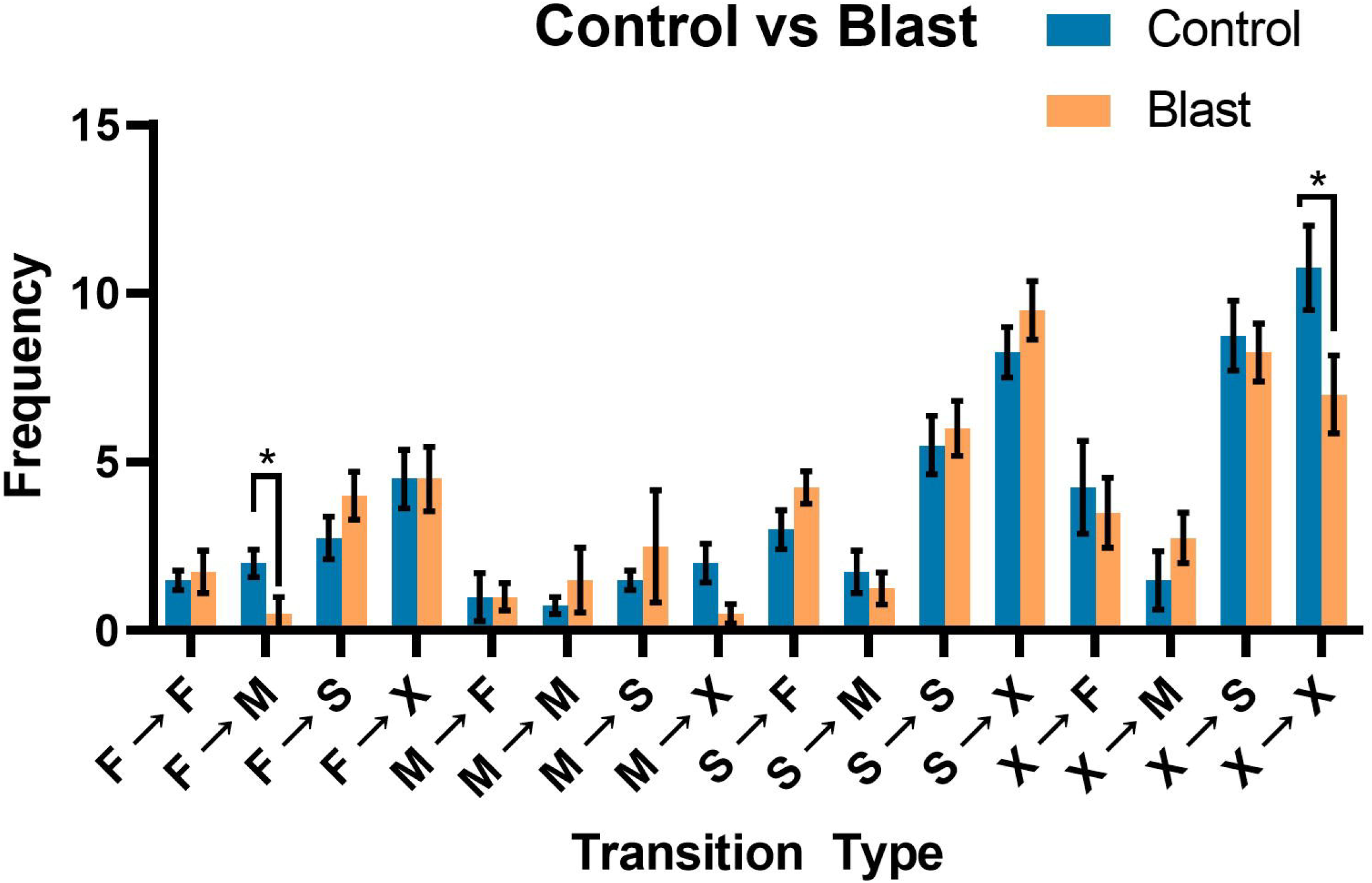
Transition states between control and blast. Quantification of network transitions between control and blast conditions. All data are presented as mean ± SEM, with n = 4 per group. Exact p-values from ANOVA with multiple comparisons are reported, with asterisks (*) denoting statistically significant differences.

**Figure 8.**
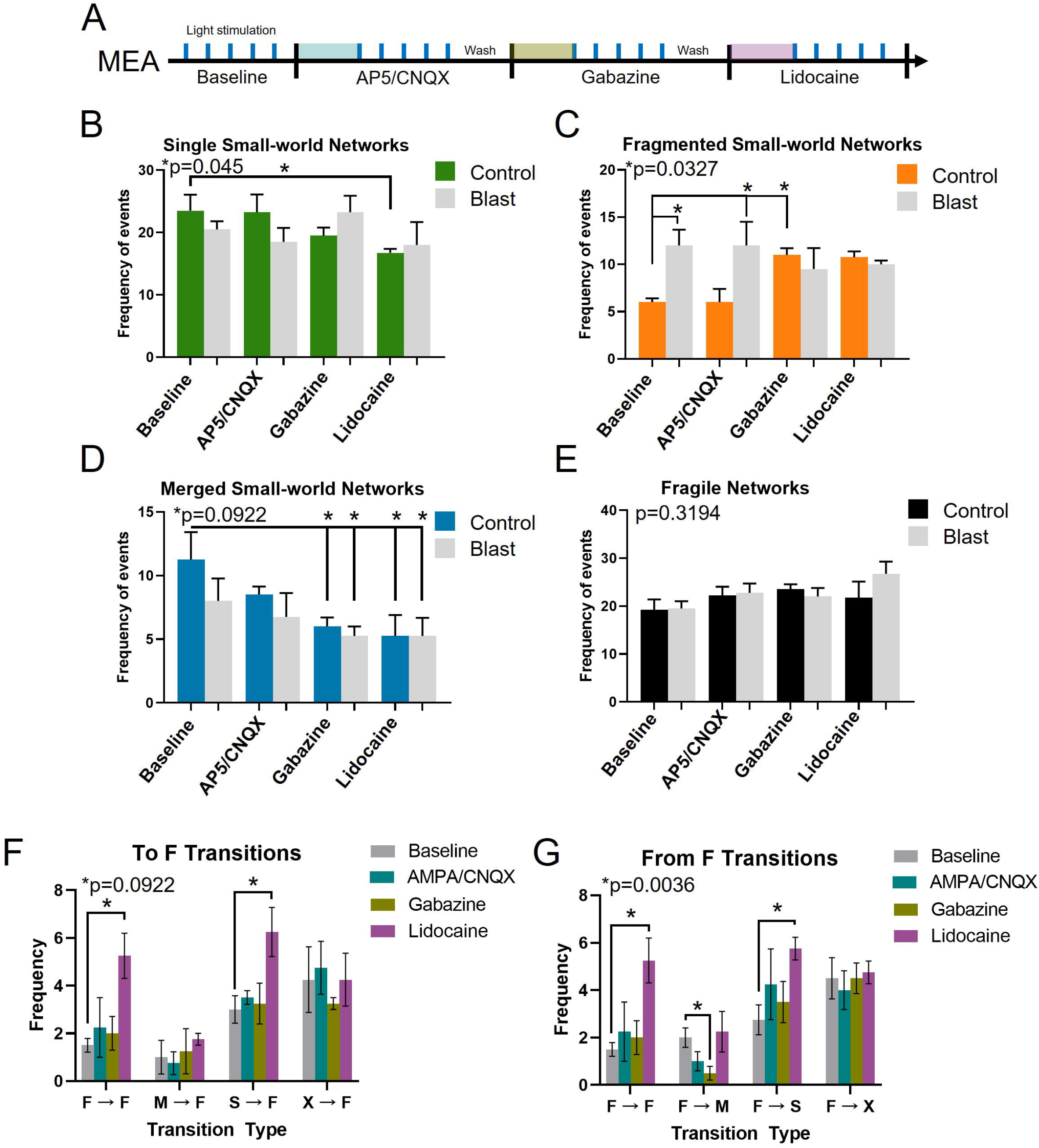
GABAergic regulation of network stability and fragmentation. (A) Schematic overview of timeline for synaptic inhibition. A baseline is recorded with pulsed blue light stimulation. A wash in period for each synaptic inhibitor precedes a stimulation test period. (B) Quantification of the frequency of single small-world networks between control and blast groups. (B) Quantification of the frequency of fragmented small-world networks. (D) Quantification of the frequency of merged small-world networks. (E) Quantification of the frequency of merged small-world networks. (F) Histogram quantifying the different network state transitions to the fragile state in response to each synaptic inhibitor. (G) Histogram quantifying the network state transitions from the fragile state. All data are presented as mean ± SEM, with n = 4 per group. Exact p-values from ANOVA with multiple comparisons are reported, with asterisks (*) denoting statistically significant differences.

For S-SWNs (Single Small-World Networks), only lidocaine produced a significant effect in control organoids, reducing the frequency of S-SWNs from 23.5 ± 2.598 to 16.75 ± 0.6292 (One-way ANOVA, F(7, 24) = 1.157, *p=0.045, Figure 5B). In contrast, for F-SWNs (Fragmented Small-World Networks), an increase was observed at baseline following blast overpressure (from 6.0 ± 0.4082 to 12 ± 1.683; One-way ANOVA with multiple comparisons, F(7, 24) = 2.698, *p=0.0327). This increase remained unchanged after the application of AP5/CNQX (6.00 ± 1.414 to 12 ± 2.517). However, following Gabazine application, a further increase in F-SWNs frequency was observed (6.0 ± 0.4082 to 11.0 ± 0.7071, One-way ANOVA with multiple comparisons, F(7, 24) = 2.698, *p=0.0009). This indicates that glutamatergic receptor inhibition does not significantly influence fragmented network formation, whereas GABA_A receptor inhibition facilitates network fragmentation, even in the absence of blast overpressure.

For M-SWNs (Merged Small-World Networks), a consistent decrease in the frequency of merged networks was observed following the application of all synaptic inhibitors. However, only after Gabazine treatment did we observe a significant decrease in the frequency of M-SWNs, independent of blast overpressure (Control baseline: 11.25 ± 2.175 to control gabazine: 6.0 ± 0.7071, blast gabazine: 5.25 ± 0.75, control lidocaine: 5.25 ± 1.652, blast lidocaine: 5.25 ± 1.436; One-way ANOVA with multiple comparisons F (7, 24) = 2.034, *p=0.0922, Figure 5D). No significant changes in the frequency of fragile network states were observed in response to the synaptic inhibitors (Figure 5E). Together these data suggest that GABA-mediated synaptic activity plays a dominant role in influencing network states and stability.

Next, we examined how synaptic inhibition influences transitions *into* fragmented (F) networks and *from* fragmented states to better understand the dynamics of network instability. Fragmented states are often associated with disorganization and represent a vulnerable network configuration. Examining transitions into fragmented states can provide insight into how easily stable network configurations deteriorate, while transitions from fragmented states reveal the network’s capacity to recover and reintegrate into more stable configurations.

Interestingly, the most frequent transition into an F state originates from the fragile (X) state. However, we observed that transitions from X → F are not significantly affected by synaptic inhibition (Figure 5E, 5F). This suggests that X states may be largely independent of synaptic function.

Of the inhibitors tested, lidocaine—which blocks global sodium channels—had the greatest impact on F state transitions. Lidocaine significantly increased F → F transitions (baseline: 1.5 ± 0.74 to lidocaine: 5.25 ± 0.97, one-way ANOVA, F(3, 48) = 10.54, *p<0.0001; Figure 5F), indicating a tendency for networks to remain fragmented. Additionally, lidocaine increased the frequency of S → F transitions (baseline: 3 ± 0.56 to lidocaine: 6.25 ± 1.78, one-way ANOVA, F(3, 48) = 10.54, *p<0.0001), suggesting that sodium channel inhibition promotes transitions into fragmented states.

When examining transitions from F states, we observed a significant effect of lidocaine on the network’s ability to return to S states (baseline: 3.25 ± 0.85, lidocaine: 3.75 ± 0.63). Additionally, gabazine administration led to a significant decrease in transitions from F → M states (baseline: 2.0 ± 0.41, gabazine: 0.5 ± 0.29; one-way ANOVA, F(3, 12) = 2.382, *p=0.0240), indicating that GABA__A_ receptor inhibition disrupts the network’s capacity to transition into the more stable merged state.

These data suggest that GABAergic activity is critical for promoting network integration and stability, as gabazine blocks the transition from fragmented to merged states. In contrast, lidocaine increased transitions both to fragmented states (S →F) and within fragmented states (F→ F), indicating that sodium channel blockade causes the network to remain or revert to fragmented configurations more frequently. These results highlight the differential effects of synaptic inhibition on network fragmentation, with GABAergic signaling supporting recovery from fragmentation, and sodium channel inhibition promoting persistent instability.

## Discussion

Our study demonstrated that primary blast overpressure induces significant fragmentation in small-world networks (SWNs) in cerebral organoids, leading to reduced overall network integration. Using multi-electrode arrays (MEAs), we observed a clear increase in network fragmentation and a decrease in F→M transitions, suggesting an inability of the network to transition to more stable states after blast exposure. Additionally, optogenetic stimulation partially restored network synchrony, indicating the potential for external stimuli to enhance network recovery. Importantly, we found that GABAergic signaling plays a critical role in maintaining network stability, as GABA receptor inhibition exacerbated fragmentation. These results suggest that the mechanical stress induced by blast overpressure disrupts normal network dynamics and that interventions targeting synaptic regulation could mitigate these effects.

The structural organization of neurons in cerebral organoids is critical for establishing functional networks that mimic aspects of the human brain. In our organoids, neurons are organized in ways that support the emergence of slow-wave oscillations (∼10 Hz), which likely contribute to the network state dynamics we observed. This hierarchical organization parallels the structure of large-scale brain networks, such as the default mode and attention networks, where localized circuits interact with broader network structures to enable adaptive and flexible behavior (Neuner et al., 2014; Kluger and Gross, 2021). The structural integrity of these neuronal circuits appears to be a key factor in maintaining network function, as fragmented or disorganized networks fail to support stable transitions between network states, a phenomenon we observed following blast overpressure.

Small-world networks (SWNs) are integral to efficient brain function because they balance local clustering with short global path lengths, optimizing both specialized processing and integrated communication (Watts and Strogatz, 1998; Sporns, 2011). These networks are essential for maintaining stability across multiple brain networks, allowing for rapid information transfer while preserving robust local processing (Bassett and Sporns, 2017) The ability of SWNs to withstand certain disruptions is crucial for maintaining cognitive and functional stability, but our results show that blast overpressure significantly fragments these networks, reducing their ability to coordinate activity across different networks. This fragmentation likely leads to the cognitive and behavioral deficits seen in traumatic brain injury (TBI) patients, where the breakdown of efficient communication networks results in impaired brain function (Hillary & Grafman, 2017).

GABAergic signaling, particularly from fast-spiking parvalbumin-positive (PV) interneurons, plays a pivotal role in maintaining the balance of excitation and inhibition within small-world networks. Through fast, precise inhibitory control, PV interneurons help synchronize neural oscillations such as gamma waves, which are essential for coordinating activity across distant regions of the brain. This inhibitory control is necessary to maintain the small-world balance between localized processing and global integration, preventing runaway excitation and supporting stable network transitions (Buzsáki, 2006; Leitch, 2024). These interneurons help synchronize neuronal activity, ensuring the stability of local circuits and preventing excessive fragmentation of the network. In diseases such as epilepsy, disruptions in GABAergic signaling lead to desynchronization and fragmented network states, similar to what we observed following blast overpressure (Krishna et al., 2020). Our findings suggest that the GABAergic system is crucial for preserving the stability of SWNs, as GABA receptor inhibition in our study further exacerbated network fragmentation, pointing to the importance of inhibitory signaling in preventing network collapse.

Fragmented networks represent a transitional and unstable state within the brain’s network architecture. These networks fail to transition into more stable configurations, such as single or merged SWNs, as we observed with a reduction in F → M transitions after blast exposure. GABAergic inhibitory neurons, particularly PV and somatostatin (SST) neurons, are likely involved in this process, as they play key roles in regulating network stability and preventing fragmentation (Buzsáki, 2006). In a recent review, (Villalobos, 2024) postulates that disinhibition, a network motif regulated by GABAB receptor signaling, plays a crucial role in modulating network transitions. Disinhibitory microcircuits in the prefrontal cortex have been demonstrated to mediate conditioned fear responses, supporting the idea that GABAergic disinhibition aids in network state transitions during social behavior and emotional learning (Xu et al., 2019). Additionally, VIP-expressing interneurons in the motor cortex mediate disinhibition by inhibiting SST neurons, which enhances excitatory input and plasticity during motor activity (Lee et al., 2013). This disinhibitory mechanism supports dynamic transitions in cortical networks and emphasizes the importance of GABAergic signaling in maintaining flexible network states. Disruptions in GABAergic signaling may hinder the network’s ability to shift out of fragmented states by reducing inhibitory control, leading to persistent fragmentation and hyperexcitability. This reflects a broader disruption in synaptic coordination, where the loss of inhibitory mechanisms allows for unchecked excitatory activity, ultimately breaking down network coherence. While these studies suggest the role of GABA signaling in network dynamics, we believe our data is the first direct evidence that GABA regulates the stability of network states through fragmentation of SWNs.

The clinical implications of our findings are significant, particularly in the context of disorders like epilepsy, where network fragmentation and instability are hallmarks of the disease. In epilepsy, GABAergic dysfunction is linked to the loss of network synchronization, leading to both hyper-synchrony during seizures and desynchronization between events (Cossart et al., 2005). Our study suggests that similar mechanisms may underlie the network fragmentation observed in TBI, where the loss of GABAergic control results in unstable network states. Pharmacological agents that modulate GABAergic signaling, such as those used to treat epilepsy (e.g., gabapentin), may have potential therapeutic applications for restoring network stability in TBI (Krishna et al., 2020). Further research is needed to explore how synaptic modulation can be used to repair disrupted networks following mechanical trauma.

## Acknowledgement

We would like to acknowledge the use of AI-assisted technologies that significantly contributed to the completion of this research. Specifically, we utilized ChatGPT, a language model developed by OpenAI, for assistance in refining ideas, editing sections of the manuscript, and providing insights into complex concepts. Additionally, Google Colab was instrumental in our data analysis, data visualization, and computational tasks. This environment facilitated not only the execution of our models but also assisted in debugging and improving our code. These technologies were invaluable in enhancing our research capabilities and streamlining the analysis process.

